# Optogenetic Amplification Circuits for Light-Induced Metabolic Control

**DOI:** 10.1101/2020.12.23.424152

**Authors:** Evan M. Zhao, Makoto A. Lalwani, Jhong-Min Chen, Paulina Orillac, Jared E. Toettcher, José L. Avalos

## Abstract

Dynamic control of microbial metabolism is an effective strategy to improve chemical production in fermentations. While dynamic control is most often implemented using chemical inducers, optogenetics offers an attractive alternative due to the high tunability and reversibility afforded by light. However, a major concern of applying optogenetics in metabolic engineering is the risk of insufficient light penetration at high cell densities, especially in large bioreactors. Here, we present a new series of optogenetic circuits we call OptoAMP, which amplify the transcriptional response to blue light by as much as 21.8-fold compared to the basal circuit (OptoEXP). These circuits show as much as a 41-fold induction between dark and light conditions, efficient activation at light doses as low as ~1%, and strong homogeneous light-induction in bioreactors of at least 5L, with limited illumination at cell densities above 40 OD_600_. We demonstrate the ability of OptoAMP circuits to control engineered metabolic pathways in novel three-phase fermentations using different light schedules to control enzyme expression and improve production of lactic acid, isobutanol, and naringenin. These circuits expand the applicability of optogenetics to metabolic engineering.

## Introduction

Engineering microbes for chemical production often leads to metabolic burden that results in reduced biomass accumulation and product yield^1^. This challenge can be addressed using dynamic metabolic control, in which inducible systems decouple growth (prioritizing resources for biocatalyst accumulation) from chemical production (focusing metabolism on desired product synthesis)^2^. Such two-phase fermentations help balance distribution of cellular resources between essential processes and the pathway(s) of interest, leading to improvements in production^3^. The transition from growth to production has traditionally been mediated in yeast using changes in carbon sources or nutrients, including galactose^4^ and methionine^5^, or by the addition of small molecule-based inducers such as doxycycline^6^. Recently, alternative approaches such as quorum sensing^7^ and optogenetics^8–10^ have emerged as new modalities of dynamic control that provide improved autonomous and user-mediated control, respectively.

Optogenetic control of gene expression involves the manipulation of photosensitive protein domains that respond to specific wavelengths of light. Compared to chemical induction, light is highly tunable via duty cycle or intensity, does not require changing medium composition, and can be flexibly and reversibly applied in any light schedule during growth and production. Many optogenetic systems have been developed in a variety of model organisms for different applications, which are reviewed elsewhere^11^. In particular, our group has applied the EL222 transcription factor^12^ from *Erythrobacter litoralis* HTCC2594 to yeast metabolic engineering^8,9^. EL222 is comprised of an N-terminal light-oxygen-voltage (LOV) domain, as well as a C-terminal helix-turn-helix (HTH) DNA binding domain. When the LOV domain absorbs blue light through flavin mononucleotide acting as a chromophore, it undergoes a conformational, shift that exposes the HTH domain, allowing it to bind to a cognate DNA sequence (C20)^13^. In darkness, EL222 spontaneously reverts to the inactivated state (half-life of ~30s^14^). Fusing the viral VP16 transactivation domain to EL222 and adding tandem repeats of C20 upstream of a minimal promoter (P_C120_) enables blue light-inducible gene expression^14^, a system which we have named OptoEXP when used in yeast^8^. Transcriptional systems such as OptoEXP can be rewired using biological logic gates to build gene circuits, raising the possibility of modifying and improving the blue light response of VP16-EL222 and P_C120_.

Optogenetic circuits have been previously designed to invert the transcriptional response to light and induce gene expression in darkness for metabolic engineering applications^8–10^. Such gene circuits use light to induce expression of a transcriptional repressor, thus inhibiting transcription of metabolic pathways when cells are exposed to light and activating them in the dark. Our group has used this approach to construct optogenetic inverter circuits that harness the GAL regulon of *Saccharomyces cerevisiae* (OptoINVRT circuits^8,9^). The GAL regulon is controlled by the Gal4p transcription factor, Gal80p repressor, and Gal3p receptor to selectively transcribe genes from the Leloir pathway for galactose metabolism^15^. OptoINVRT circuits use OptoEXP to control the expression of *GAL80*, thereby inhibiting Gal4p in the light and making Gal4p-activated promoters darkness-inducible ^8,9^. These inverter circuits were originally designed to address potential light penetration challenges in microbial fermentations, reasoning that inducing the production pathways with darkness would largely avoid the high cell densities that limit light penetration. Consequently, the application of optogenetics to metabolic engineering has thus far been limited to inducing metabolic pathways in darkness. However, the ability to efficiently, induce metabolic with light in addition to darkness would greatly expand the potential of optogenetics for metabolic engineering applications.

In this study, we present blue light-activated “OptoAMP” circuits that reach high levels of gene expression with minimal light exposure in *S. cerevisiae* by amplifying the transcriptional response to light. These circuits amplify the optogenetic signal of OptoEXP using the GAL regulon, allowing control over a suite of Gal4p-activated promoters with different light activation strengths and sensitivities. Furthermore, we increase the light sensitivity of our circuits by incorporating mutations to the EL222 protein that modify its deactivation kinetics, achieving robust light-dependent gene expression at high cell densities in lab-scale bioreactors even when exposed to light only 5% of the time. OptoAMP circuits enable optimization of chemical production by selectively applying light during different stages of fermentation, leading to improved production of lactic acid, isobutanol, and naringenin. Our results establish light (in addition to darkness) as a powerful induction tool for microbial chemical production, even in cases where light penetration may be limited.

## Results and Discussion

### “OptoAMP” circuits amplify optogenetic signals by controlling GAL4 expression

Lowering the light requirements to activate optogenetic circuits would allow light-induced gene expression in larger bioreactors and at the high cell densities typically encountered during the production phase of chemical fermentations. To achieve this, we exploited the GAL regulon of *Saccharomyces cerevisiae* (Fig. 1a). After deleting the endogenous copies of *GAL4* and *GAL80*, we integrated *GAL4* under the P_C120_ promoter controlled by VP16-EL222, and a copy of constitutively expressed VP16-EL222 (P_TEF_), resulting in OptoAMP1. To test this circuit, we also co-integrated GFP expressed using various promoters natively activated by Gal4p: P_GAL1_, P_GAL10,_ P_GAL2_, and P_GAL7_. These strains (see Supplementary Tables 1-2) show blue light-dependent GFP expression, with different promoters showing varying light sensitivities and fold-changes in maximum expression relative to OptoEXP (Fig. 1b). P_GAL1_ and P_GAL7_ express 80% and 76% (respectively) of maximal expression when exposed to 10% light dose (compared to 57% for OptoEXP), while P_GAL2_ and P_GAL10_ show full activation when exposed to 10% light dose. Additionally, while OptoAMP1 can amplify the transcriptional response of OptoEXP by more than 7-fold (when controlling P_GAL1_), none of these *GAL* promoters reach the expression levels achieved by the strong constitutive promoters P_TDH3_ and P_TEF1_, with the strongest P_GAL1_ promoter reaching 73% of P_TEF1_ expression in full blue light. Therefore, additional engineering is required to attain the promoter strengths often necessary for chemical production.

**Figure 1.**
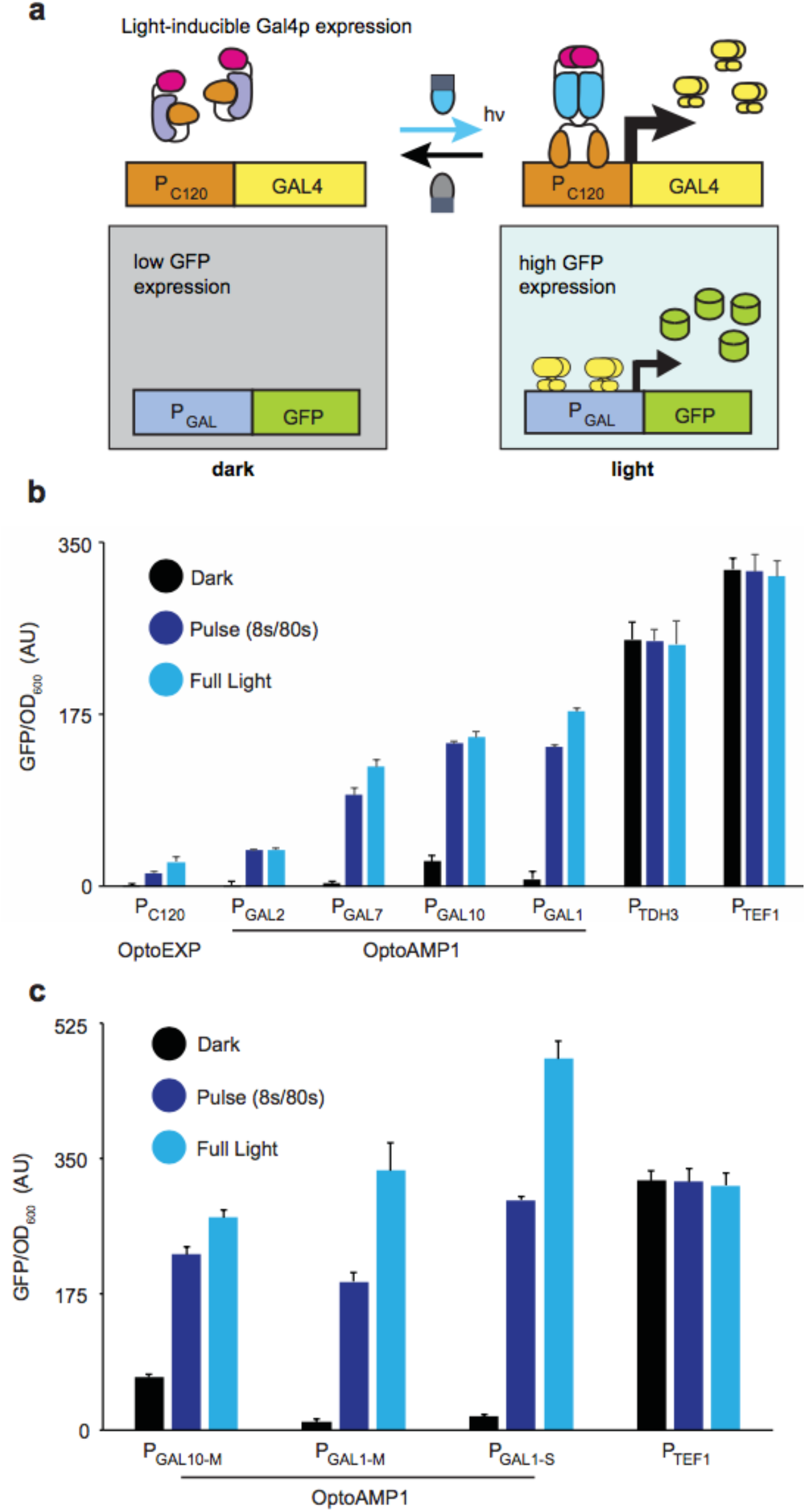
Characterization of OptoAMP1 circuit and promoters. (a) OptoAMP1 amplifies the transcriptional activity of VP16-EL222 in blue light by expressing *GAL4* from the P_C120_ promoter using OptoEXP, which then drives higher levels of gene expression from *GAL4*-activated promoters. (b) GFP expression with OptoEXP and different Gal4p-activated promoters controlled by OptoAMP1 under different light conditions: OptoEXP (YEZ139), P_GAL2_ (YEZ142), P_GAL7_ (YEZ143), P_GAL10_ (YEZ141), P_GAL1_ (YEZ72). Intermediate pulsing was performed using an 8s ON / 72s OFF (10%) light duty cycle. (c) GFP expression with engineered promoters controlled by OptoAMP1 under different light conditions: P_GAL10-M_ (YEZ189), P_GAL1-M_ (YEZ133), P_GAL1-S_ (YEZ214). Intermediate pulsing was performed using an 8s ON / 72s OFF (10% light dose) duty cycle. AU, arbitrary units of fluorescence and optical density. All data are shown as mean values; error bars represent the s.d. of four biologically independent 1-ml sample replicates exposed to the same conditions.

Engineering promoters to increase maximum expression level provides an opportunity to further increase amplification capabilities^16^. To develop a set of strong light-inducible promoters, we removed the binding sites of Mig1p, a global repressor that binds in the presence of glucose, within P_GAL10_ and P_GAL1_^9^. These mutants, P_GAL10-M_ and P_GAL1-M_, increase GFP expression levels by 39% and 45%, respectively, compared to the wild-type promoters (Fig. 1c). Addition of a P_GAL1_ fragment containing four additional Gal4p binding sites upstream of the P_GAL1-M_ promoter, results in a super-enhanced *GAL1* promoter named P ^9^, which further enhances expression by 43%. P_GAL1-S_ reaches an activation level under full blue light that is 52% higher than that of P_TEF1_, a strong constitutive promoter commonly used in metabolic engineering. Although these promoters are tunable by varying the light duty cycle, they still require at least 10% light dose to reach maximal gene expression levels, leaving ample room for improvement.

### EL222 variant with prolonged lit-state activation enhances circuit sensitivity

To reduce the amount of light required to activate our optogenetic circuits, we altered the photosensitivity of VP16-EL222. Previous studies have shown that EL222 mutations affect its lit- or dark-state half-life and transition kinetics^13,17^. We made the A79Q substitution in EL222, reported to increase its lit-state half-life from 30 seconds to 300 seconds^13^, which when used with a VP16 fusion to activate P_C120_ results in a new circuit we call OptoEXP2. Using GFP to measure gene expression under different light duty cycles, we found that OptoEXP2 achieves 78% of maximal activation with only 2 seconds of illumination in 80-second periods, and has 3.5-fold stronger maximal expression than OptoEXP under full light (Supplementary Fig. 1a,b). However, OptoEXP2 is also ~31 times leakier in the dark than OptoEXP. Increasing the active-state half-life of VP16-EL222 not only increases light sensitivity, but also raises maximum expression level, likely due to longer binding of VP16-EL222 to P_C120_. We combined this VP16-EL222 (A79Q) variant with *GAL4* expression from P_C120_ to create OptoAMP2. When OptoAMP2 is used to control GFP expression from P_GAL1-S_, the maximum GFP level achieved (under full blue light) is 2.6-fold higher than what is obtained with P_TEF1_. However, this circuit is very leaky, exhibiting 48% of the expression from P_TEF1_ even in complete darkness, (Supplementary Fig. 1b). Therefore, while OptoEXP2 and OptoAMP2 are significantly more light-sensitive than OptoEXP1 and OptoAMP1, respectively, additional modifications are necessary to restore a tight OFF-state repression.

Effective dynamic control in metabolic engineering requires that the enzymes under control have a low background level of gene expression. We previously showed that controlling protein stability of gene circuit components with photosensitive degradation domains can reduce leakiness^8,9^. We expressed the Gal80p repressor fused to a C-terminal photo-sensitive degron domain (PSD), reasoning that it would repress leaky Gal4p activity in the dark, but also get actively degraded in blue light to allow Gal4p-mediated transcriptional activation (Fig. 2a). We varied the expression level of the Gal80p-PSD fusion using two moderate-strength constitutive promoters^18^, P_ADH1_ and P_RNR2_, resulting in OptoAMP3 and OptoAMP4, respectively. Both circuits are tunable using light doses as low as 0.8% (1s ON / 119s OFF) and show low leakiness in the dark (2.5% and 3.2% of maximal activity for OptoAMP3 and OptoAMP4, respectively). Both circuits show strong light response amplification: for example, OptoAMP3 requires only a 1.7% light duty cycle to reach 52% of its maximum expression, which corresponds to 116% the expression level of P_TEF1_. Furthermore, the maximal activity of OptoAMP4 (full light) is 15% higher than that of OptoAMP3, corresponding to 211% of P_TEF1_ activity. The higher strength of OptoAMP4 is likely due to the weaker promoter (P_RNR2_) used to express *GAL80-PSD* in this circuit, relatively to OptoAMP3 (P_ADH1_)^18^. However, OptoAMP3 has a higher light-to-dark fold change in activity (40.7 vs 30.8 of OptoAMP4), which makes both circuits potentially useful for different applications.

**Figure 2.**
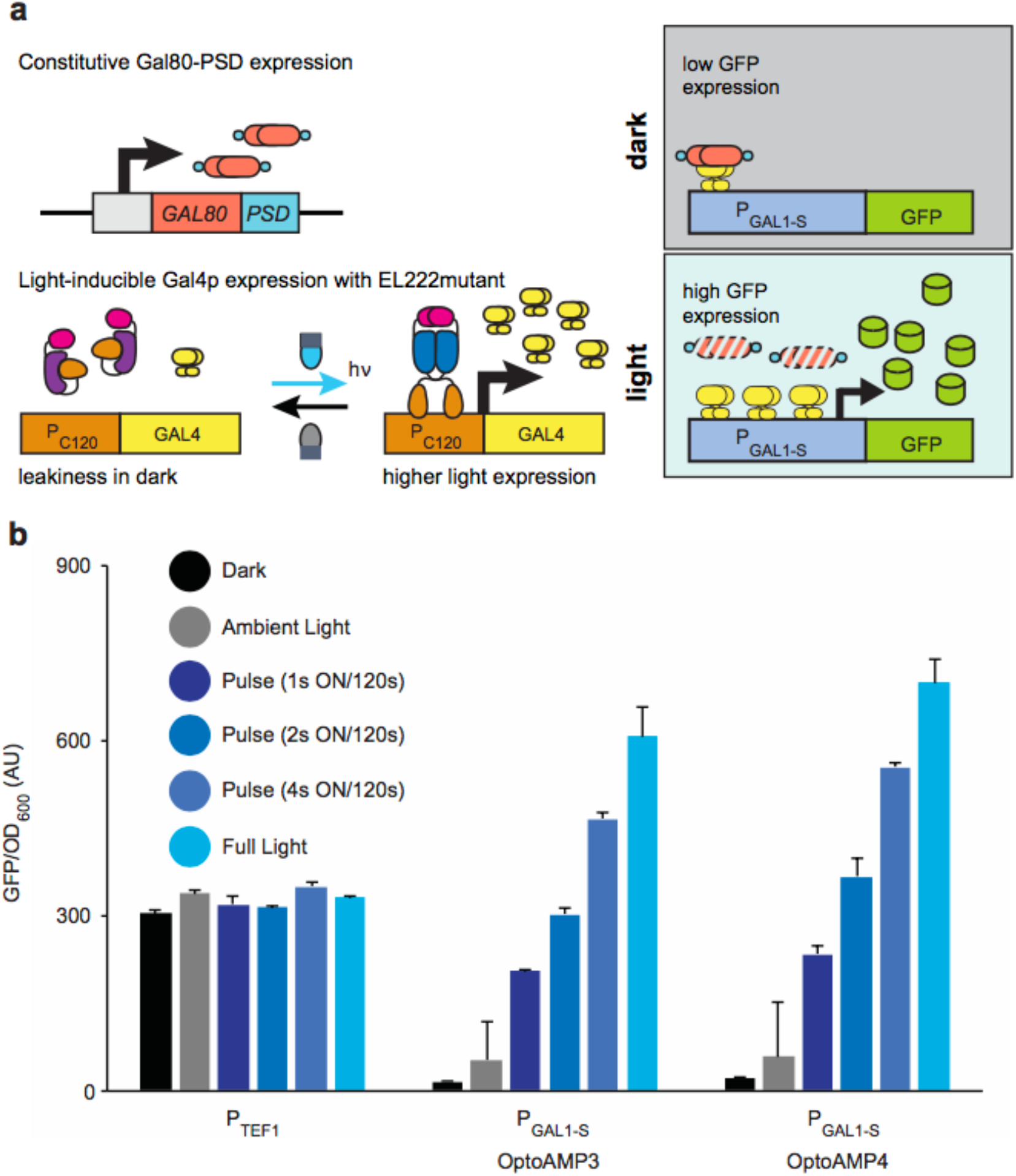
Characterization of OptoAMP3 and OptoAMP4. (a) Expressing Gal80p C-terminally tagged with a photosensitive degron domain reduces leaky activity of Gal4p in the dark, without repressing Gal4p activity in blue light. (b) GFP expression using P_GAL1-S_ controlled by OptoAMP3 (YEZ337) or OptoAMP4 (YEZ336) under different light conditions, compared to constitutive P_TEF1_. Intermediate light pulsing was performed using 1s ON/119s OFF (0.83%), 2s ON/118s OFF (1.7%), and 4s ON/116s OFF (3.3%) duty cycles. AU, arbitrary units of fluorescence and optical density. All data are shown as mean values; error bars represent the s.d. of four biologically independent 1-ml sample replicates exposed to the same conditions.

### OptoAMP4 facilitates light-induced gene expression at high cell density

OptoAMP circuits are designed to overcome light-limited conditions in bioreactors (at least lab-scale) at high cell densities. To demonstrate their enhanced functionality, we compared the ability of OptoAMP4 to induce gene expression with light in a 5L bioreactor at high cell density against that of our previous OptoEXP circuit^8^. We inoculated 3L of media containing 15% glucose (in a 5L bioreactor) with strains using OptoEXP or OptoAMP4 to induce GFP expression with light. Starting from an initial OD_600_ of 0.1, we grew the culture in darkness until it reached an OD_600_ of 10. We then exposed only 7% of the culture’s bulk surface (through one side of the partially covered glass reactor vessel using blue LEDs) to a light duty cycle of 5s ON / 95s OFF (Fig. 3a, See Methods). Under these conditions, OptoEXP shows no detectable light-induced GFP expression after 24 hours of starting the light exposure (Fig. 3b). In contrast, OptoAMP4 shows homogeneous expression of GFP after only three hours of limited light induction (Fig. 3c). Furthermore, OptoAMP4 maintains its GFP expression even after 24 hours of batch fermentation, at which point the OD_600_ of the culture is 41.2 – well into stationary phase. These results demonstrate the feasibility of using OptoAMP4 for light-induced gene expression in fermentations at relatively high cell densities, in bioreactors of at least 5L, and using low light doses, which are conditions with substantially limited light penetration that prevent activation of earlier optogenetic circuits (OptoEXP).

**Figure 3.**
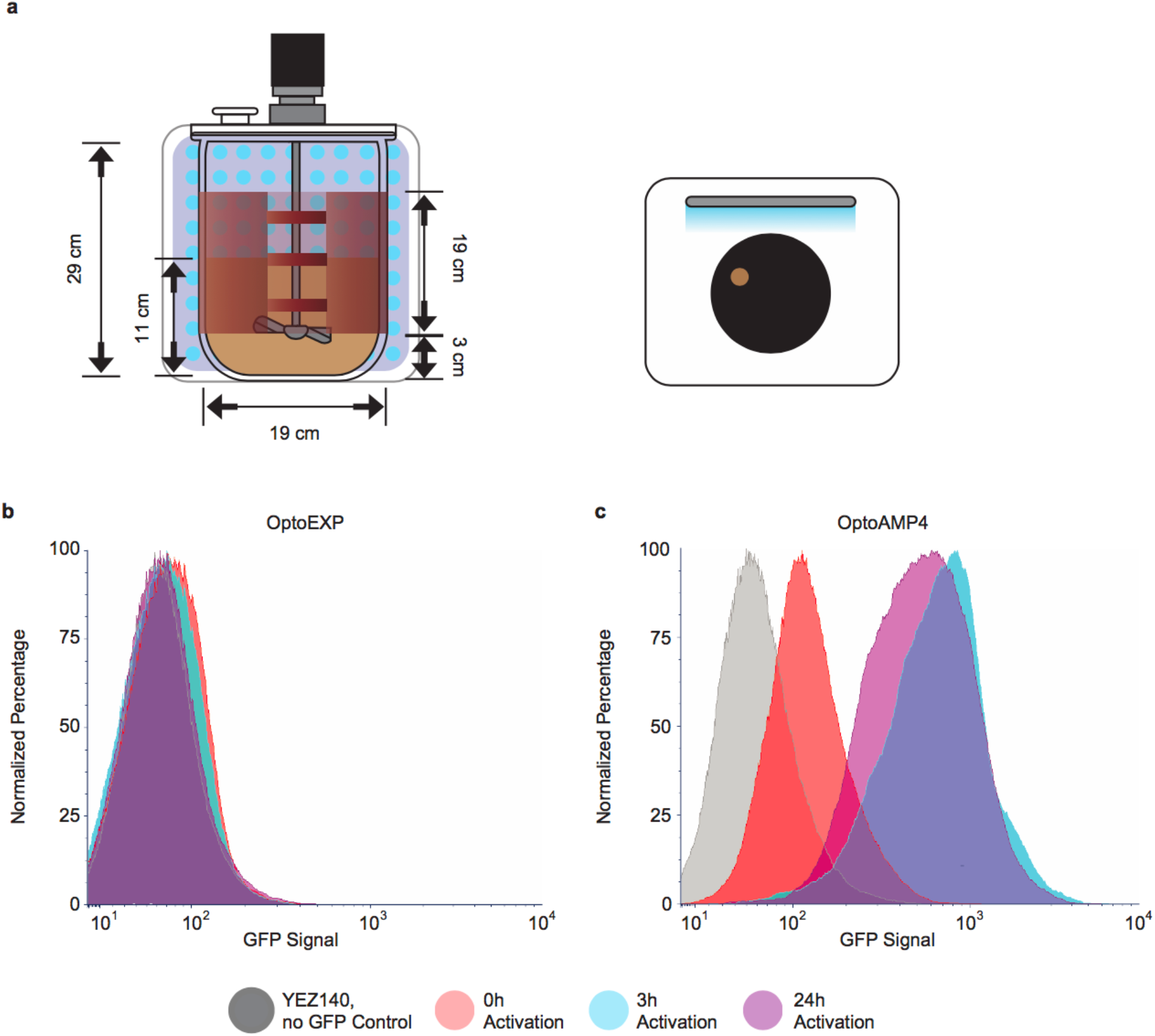
GFP expression in laboratory-scale fermenter with controlled light exposure. (a) Schematic of 5-L fermentor setup with the dimensions of the area exposed to light. Red is the heating blanket around the reactor. Tan color depicts the cell culture (3 L). (b,c) Representative flow cytometry results using YEZ139, which has OptoEXP controlling GFP expression from P_C120_ (b) or YEZ336, which has OptoAMP4 controlling GFP expression from P_GAL1-S_ (c). Cells were grown in the dark in batch using 15% glucose. When cultures reached an OD_600_ of ~15, they were exposed to 5s ON / 95s OFF illumination for the rest of the experiment. Red histograms show fluorescence levels at pre-induction at an OD_600_ of ~15; blue is after 3 h of induction at an OD_600_ of ~23 (b) or ~22 (c); Purple is after 24 h of induction at an OD_600_ of ~41. Grey is strain YEZ140 without GFP, which was used as a negative control. Each curve is generated from 10,000 single-cell events.

### OptoAMP circuits enhance production of valuable chemicals

Having established that OptoAMP circuits can overcome limitations in light penetration at least in lab scale bioreactors, we sought to determine whether they could be used to control biosynthetic pathways with minimal light stimulation. A unique benefit of using optogenetics to control microbial fermentations is that distinct light schedules may be applied to design any number of fermentation protocols. Thus, we developed a new fermentation protocol consisting of three phases: i) a growth phase, in which biomass accumulates; ii) an induction phase, in which OptoAMP activates production pathways; and iii) a production phase, in which strains produce chemicals of interest after resuspension in fresh media under variable light conditions (Fig. 4a, Supplementary Figure 2). With this new fermentation protocol, we can apply various light conditions (full light, full darkness, or light pulses) in each phase, to maximize production.

**Figure 4.**
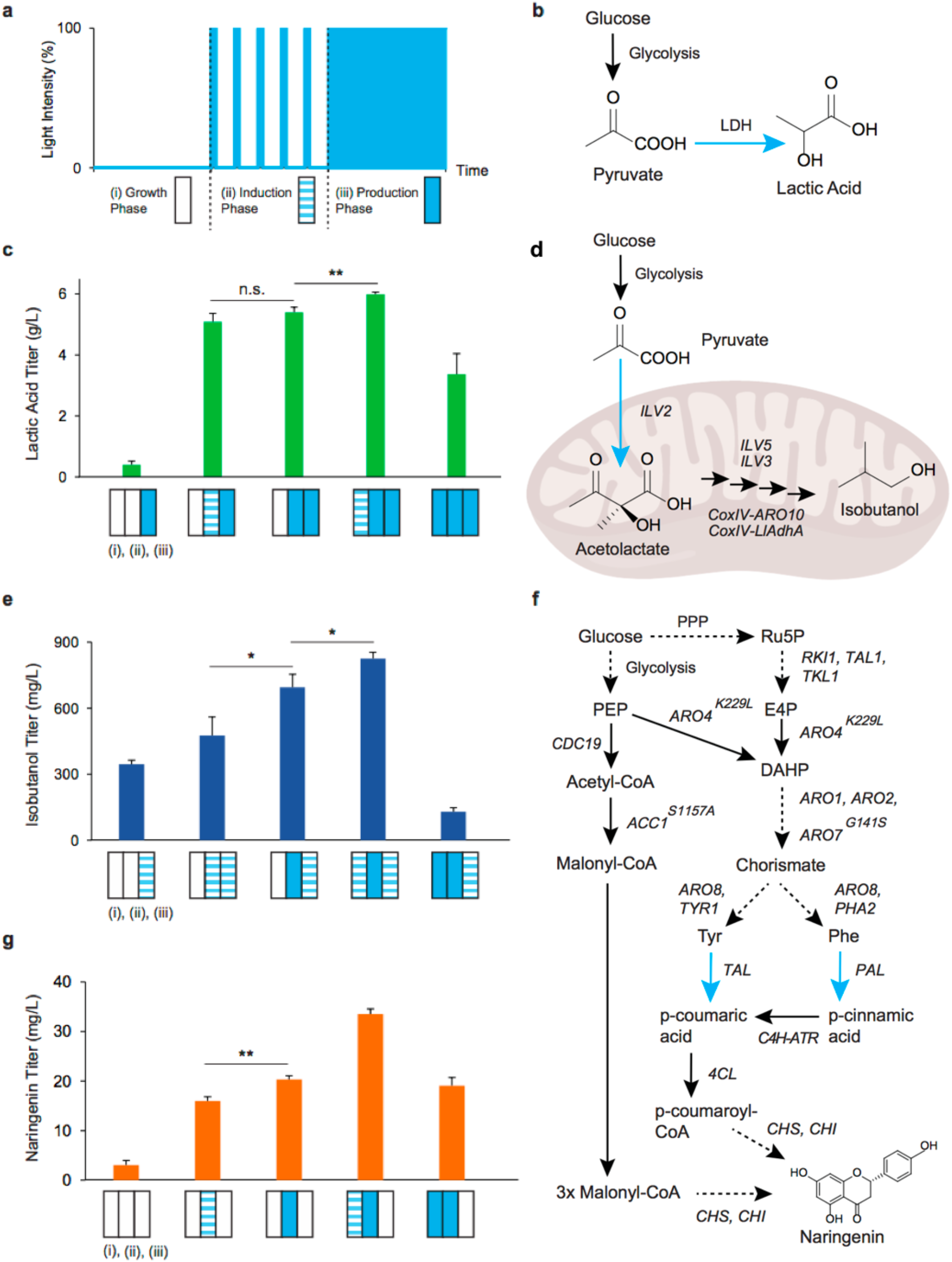
Chemical production OptoAMP4-mediated light induced 3-phase fermentations. (a) Example of a light schedule on a 3-phase fermentation, showing a growth phase (i) in full darkness, an induction phase (ii) with a representative light pulse, and a production phase (iii) in full blue light. (b) Lactic acid biosynthesis, with optogenetic control of LDH (blue arrow). (c) Lactic acid production using OptoAMP4 to control LDH expression from P_GAL1-S_ (YEZ423) using different light schedules in the growth and induction phases, each phase lasting: (i) growth, 20h, (ii) induction, 12h, and (iii) production, 24h. (d) Mitochondrial isobutanol production pathway, with *ILV2* expression controlled optogenetically (blue arrow). Mitochondrion was created using Biorender.com. (e) Isobutanol production using OptoAMP4 to control *ILV2* expression from P_GAL1-M_ (YEZ516) using different light schedules in the growth and induction phases, each phase lasting: (i) growth, 20h, (ii) induction, 12h, and (iii) production, 48h. (f) Naringenin biosynthetic pathway, with FjTAL and AtPAL2 expression controlled optogenetically (blue arrow). Multi-enzymatic steps are shown in dashed arrows. (g) Naringenin production using OptoAMP4 to control the expression of FjTAL and AtPAL2 from P_GAL1-S_ (YEZ488) using different light schedules in the growth and induction phases, each phase lasting: (i) growth, 20h, (ii) induction, 12h, and (iii) production,48h. \**P* < 0.05, \*\**P* < 0.01. Statistics are derived using a two-sided *t*-test. All data are shown as mean values; error bars represent the s.d. of four biologically independent 1-ml sample replicates exposed to the same conditions.

We first tested our new fermentation protocols to test OptoAMP4 in the production of lactic acid (LA), a valuable polymer precursor and food additive that only requires one exogenous enzyme (lactate dehydrogenase, LDH) to synthesize from pyruvate (Fig. 4b). We used OptoAMP4 driving P_GAL1-S_ to control LDH expression during our three-phase fermentation protocol. We applied different light schedules (dark, full light, or 1s ON / 79s OFF light pulses) during the growth (20h) and induction (12h) phases, but always full light during the production (24h) phase to maximize LDH induction. Applying only 1.3% light pulses during the induction phase leads to a 12.8-fold increase in LA titer (5.1 ± 0.3 g/L), relative to keeping cells in the dark during the induction phase (0.4 ± 0.1 g/L) (Fig. 4c). Furthermore, there is a statistically significant improvement in LA production (6.0 ± 0.1 g/L) when a 1.3% light pulse is applied during the growth phase, compared to growing the cells in complete darkness (5.4 ± 0.2 g/L) or keeping the light on throughout the three phases of fermentation (3.4 ± 0.7 g/L). These results suggest that chemical production may be improved by weak early pathway induction in new multiphasic fermentation protocols, a unique capability of optogenetic controls, enabled by the enhanced light sensitivity of OptoAMP circuits.

We next developed a strain with light-induced isobutanol production to show that OptoAMP circuits can regulate production of a multi-step biosynthetic pathway with low levels of light exposure during the production phase. Using OptoAMP4, we controlled the expression of the first gene of the mitochondrial isobutanol pathway (*ILV2*)^19^ from P_GAL1-M_ with light, while the rest of the genes in the pathway^19^ (*ILV5*, *ILV3*, CoxIV-*ARO10*, and CoxIV-*adhA^RE1^* from *Lactococcus lactis*) were constitutively expressed (Fig. 4d). We ran three-phase light-induced isobutanol fermentations, in which the cell cultures were exposed to different light schedules, (dark, full light, or 1s ON / 79s OFF light pulses) during the growth (20h) and induction (12h) phases, but always exposed to a limited 1.7 % light dose (2s ON / 118s OFF) during the production phase (48h) to mimic the limited light penetration conditions typically found in larger bioreactors. Isobutanol titers reach 350 ± 20 mg/L even without light exposure until the production phase, probably because the endogenous copy of *ILV2* is still present in the strain background (Fig. 4e). However, applying 1.3% light or full light during the induction phase leads to a 1.4-fold or 2-fold improvement in titer (480 ± 80 and 700 ± 60 mg/L), respectively, consistent with light-induced pathway expression prior to the production phase. Applying a low light dose during the growth phase leads to a further 19% increase in isobutanol titers (830 ± 30) compared to keeping the cultures in the dark during the same phase (690 ± 60 mg/L). In contrast, exposing cultures to full light during the growth phase leads to 6.4-fold lower isobutanol titer (130 ± 20 vs. 830 ± 30 mg/L). These results again imply that early (but not excessive) pathway induction during the growth phase can improve chemical production, similar to what we observed for LA production.

Some metabolic pathways may benefit from more complex fermentation protocols afforded by the ease with which light can be instantly applied and removed. Such is the case for the biosynthesis of naringenin, a flavonoid with anti-inflammatory and other therapeutic properties derived from tyrosine and phenylalanine (Fig. 4f). Starting from a strain background that is engineered to upregulate aromatic amino acid production^20,21^ (see Methods), we introduced enzymes of the naringenin pathway^22^ (AtC4H-AtATR2, At4CL2, HaCHS, and PhCHI) under constitutive promoters (see Supplementary Table 2). In addition, we used OptoAMP4 controlling P_GAL1-S_ to drive expression of the first enzymatic steps to produce naringenin from tyrosine and phenylalanine: tyrosine ammonia-lyase (FjTAL) and phenylalanine ammonia-lyase (AtPAL) (Fig. 4f). We carried out three-phase fermentations for naringenin production, testing various light conditions in all three phases. As with LA and isobutanol, naringenin production benefits from a low light dose (1.3%) during the growth phase (20h), achieving 76% and 65% higher titers than if this phase is kept in full light or darkness, respectively (Figure 4g). Moreover, naringenin titers improve 6.8-fold when the production phase (48h) is kept in complete darkness compared to under full light (Supplementary Figure 3). This suggests that the levels of FjTAL and AtPAL accumulated during the induction phase (12h) are sufficient to achieve higher naringenin production, making light exposure during the production phase unnecessary and perhaps detrimental due to cellular overburden or photosensitivity of the product or its precursors. Optogenetic control of naringenin production is thus a good example of how specifically tailored light schedules in three-phase designer fermentations may benefit different metabolic pathways.

## Conclusions

Optogenetics has the potential to change control of microbial chemical production through flexible and tunable induction of target enzymes. The first optogenetic circuits developed for metabolic engineering employed inverter circuits that induce biosynthetic pathways of interest with darkness^8–10^. This preemptive design principle was used to circumvent potential limitations in light penetration during the production phase of fermentations, when cell densities are typically at their highest. However, to realize the full potential of optogenetics to control, automate, and optimize microbial fermentations for chemical production, we must have the ability to induce gene expression with light as effectively as with darkness. This would allow the design of new complex fermentation protocols with more phases than the biphasic growth/production protocols typically used. In addition, it would make it possible to use periodic light pulse operations^8^ in larger bioreactors and at higher cell densities.

In this study, we present a blueprint for amplifying blue light-activated signals that balances maximum expression strength, light sensitivity, and tight OFF-state control. The suite of Gal4p-activated promoters, which we have made light-inducible through OptoAMP circuits (Fig. 1b), provides the promoter diversity required for controlling longer metabolic pathways by offering different strengths for tuning pathway expression levels, as well as avoiding tandem repeats that decrease strain stability. Furthermore, the compatibility of our circuits with media containing glucose is an added benefit over traditional galactose-based induction, avoiding the need for media changes and simplifying process operation. Such features make our OptoAMP circuits powerful additions to the growing yeast optogenetic toolkit.

OptoAMP circuits, enabled by a hypersensitive photosensory domain variant of EL222 and the transcriptional strength of Gal4p, can amplify the transcriptional response to light by as much as 21.8-fold, relative to our original OptoEXP circuit^8^. To simulate the limited light penetration conditions that would be found in large-scale bioreactors at high cell densities, we reduced the light-exposed bulk surface in our 5L bioreactor to 7% and the light dose applied to 5% (5s ON / 95s OFF). The robust light-induced expression attainable at OD_600_ > 40 using OptoAMP4, even under limited light illumination in lab-scale bioreactors (Fig. 3), suggests this circuit could enable the use of optogenetics to control much larger scale bioreactors using higher light doses and larger areas of exposure.

The ability to induce gene expression effectively with light, provided by OptoAMP circuits, facilitates the development of complex fermentation protocols that include illumination of high cell density cultures to induce biosynthetic pathways of interest. We demonstrate this capability by developing a new three-phase fermentation protocol, which we tested in the production of three different valuable chemicals (LA, isobutanol, and naringenin). We found that while high light doses during the induction phase improve production, early (but not excessive) light exposure during the growth phase is also important to increase titers of all three chemicals. Naringenin production is a particularly interesting case, in which a dark production phase is most effective, probably due to photosensitivity of the product and precursors^23^. This result shows that OptoAMP4 is strong enough to induce and accumulate sufficient enzymes (and mRNA) during the induction phase, which improve naringenin titers even in a dark production phase. The fact that different light schedules could be found to maximize production of each chemical, points to the huge potential of using optogenetics and complex fermentation protocols to optimize chemical production.

By reducing the light requirements for homogeneous gene expression through optogenetic circuit design, OptoAMP circuits add a promising new layer of control over microbial fermentations. Using carefully timed light pulses over multiphasic fermentation protocols, it may be possible to optimize complex metabolic pathways in unprecedented ways. We anticipate that our optogenetic amplifiers, combined with optogenetic inverters and multichromatic systems, will enable more elaborate metabolic control in the future, where light as well as darkness can serve as strong inducing agents. These OptoAMP circuits thus constitute an important milestone, towards the future application of better optogenetic controls in larger, even industrial-scale, bioreactors, where light penetration is limited.

## Methods

### Assembly of DNA constructs

We cloned promoter-gene-terminator sequences into previously described standardized vector series (pJLA vectors) as previously described^19^ (Supplementary Figure 4). Plasmids were transformed into chemically competent *Escherichia coli* strain DH5α. Qiagen Miniprep, Qiagen Gel Extraction, and Qiagen PCR purification kits were used to extract and purify plasmids and DNA fragments, following manufacturer’s instructions. The EL222 mutant was ordered as a gBlock from Integrated DNA Technologies. All vectors were sequenced with Sanger Sequencing from GENEWIZ before using them to transform yeast. All plasmids used in this study are catalogued in Supplementary Table 1.

### Yeast strains and transformations

Yeast transformations were carried out using standard lithium acetate protocols^24^, and the resulting strains are catalogued in Supplementary Table 2. Gene constructs derived from pYZ12-B, pYZ162, and pYZ23 vectors were genomically integrated into the *HIS3* locus, *LEU2* locus, or δ-sites (YARCdelta5), respectively. Zeocin (Thermo Fisher Scientific), ranging from 800 to 1200 μg/mL, was used to select for strains with δ-integrations. Gene deletions were carried out by homologous recombination as previously described^8^. All strains with gene deletions were genotyped with PCR to confirm their accuracy. We integrated constructs into the *HIS3* locus, *LEU2* locus, or δ-sites to promote strain stability (Supplementary Table 2). We avoid using, tandem repeats to prevent recombination after yeast transformation, and thus do not observe strain instability.

### Yeast cell culture growth, centrifugation, and optical measurements

Liquid yeast cultures were inoculated from single colonies and grown in 24-well plates (USA Scientific Item #CC7672-7524), at 30°C and shaken at 200 rpm (19 mm orbital diameter), in either YPD or SC-dropout media with 2% glucose. To stimulate cells with light, we used blue LED panels (HQRP New Square 12” Grow Light Blue LED 14W), placed 40 cm from cell cultures. At this distance, the light panel outputs ranged from 52 μmoles/m^2^/s to 94 μmoles/m^2^/s, measured with a quantum meter (Model MQ-510 from Apogee Instruments). Light duty cycles were set using a Nearpow Multifunctional Infinite Loop Programmable Plug-in Digital Timer Switch to control the LED panels. Cell cultures were centrifuged (Sorvall Legend XTR) at 1000 RPM for 10 minutes using 24-well plate rotor adaptors.

Fluorescence and optical density (OD_600_) measurements were taken using a TECAN plate reader (infinite M200PRO). The excitation and emission wavelengths used for GFP fluorescence measurements were 485 nm and 535 nm, respectively. To process fluorescence data, the background fluorescence from the media exposed to the same light conditions was first subtracted from measured values. Then, the GFP/OD_600_ values of cells lacking a GFP construct but exposed to the same light conditions were subtracted from the fluorescence values (GFP/OD_600_) of each sample to correct for light bleaching of the media and cell contents. Reported values were calculated per the following formula.

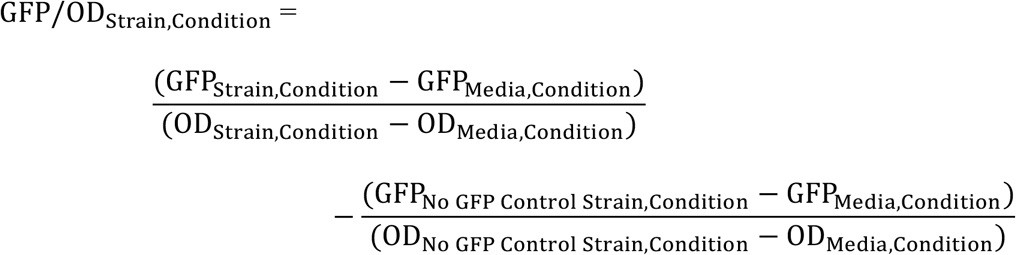

All fluorescence measurements were taken at the end of experiments or on samples taken from bioreactor cultures, such that potential activation of VP16-EL222 by the light used to excite GFP did not affect experimental progression or final results^8,9^. Controls using constitutive P_TEF1_ to express GFP showed that this fluorophore does not bleach under the light conditions tested.

To measure cell concentration, optical density measurements were taken at 600 nm wavelength, using media (exposed to the same conditions as the yeast) as blank. Measurements were done with the TECAN plate reader (infinite M200PRO) or Eppendorf spectrophotometer (BioSpectrometer basic), from samples diluted to a range of OD_600_ of 0.1 to 1.0.

### Flow Cytometry

GFP fluorescence of bioreactor experiments was quantified by flow cytometry using a BD LSR II flow cytometer and BD FacsDiva 8.0.2 software with a 488 nm laser and 525/50 nm bandpass filter. The gating used in our analyses was defined to exclude particles that are either too small or too large to be single living yeast cells (Supplementary Figure 5), based on the side scatter (SSC-A) vs forward scatter (FSC-A) plots as previously described^8^. Median fluorescence values were determined from 10,000 cells. Data were analyzed using FCS Express (De Novo Software, Pasadena, CA, USA).

### Construction and characterization of OptoAMP systems

All circuits and promoters were characterized in yeast strain YEZ44 (CENPK.2-1C, *gal80-Δ*, *gal4-Δ*). OptoAMP1 controlling GFP expression from P_GAL1_, P_GAL10_, P_GAL2_, or P_GAL7_ was integrated into the *HIS3* locus of YEZ44 to make YEZ72 (EZ-L175), YEZ141 (EZ-L319), YEZ142 (EZ-L320), and YEZ143 (EZ-L321), respectively. OptoAMP1 controlling GFP expression from P_GAL10-M_, P_GAL1-M_ and P_GAL1-S_ was integrated into the *HIS3* locus of YEZ44 to make strains YEZ189 (EZ-L380), YEZ133 (EZ-L316), and YEZ214 (EZ-L444) respectively. OptoEXP2 driving GFP expression from P_C120_ or OptoAMP2 driving GFP expression from P_GAL1-S_ were integrated into the *HIS3* locus of YEZ44 to make strains YEZ293 (EZ-L545) and YEZ292 (EZ-L560), respectively. Single colonies of each strain were inoculated into SC-His media + 2% glucose media in 24-well plates and grown overnight; cultures were kept in darkness (covered in aluminum foil) to avoid premature activation of optogenetic systems. Each culture was then back-diluted into fresh media to an OD_600_ = 0.1, and grown for 8 hours under full light, complete darkness, or light pulses of 8s ON / 72s OFF.

OptoAMP3 or OptoAMP4 driving GFP expression from P_GAL1-S_ were integrated into the *HIS3* locus of YEZ44 to make strains YEZ337 (EZ-L583) and YEZ336 (EZ-L582), respectively. We monitored GFP expression as above, exposing cells to full light, ambient light (warm room with lights turned off), complete darkness, or light pulses of 1s ON / 119s OFF, 2s ON / 118s OFF, or 4s ON / 116s OFF.

### 5L Bioreactor testing of OptoAMP4 and OptoEXP

To test OptoEXP or OptoAMP4 in higher cell density conditions, we inoculated YEZ139 (OptoEXP) or YEZ336 (OptoAMP4) into 10 mL of SC-His + 2% glucose from a single colony and grew in the dark for 16 h. We then set up a BioFlo120 system with a 5 L bioreactor (Eppendorf, B120110002) and added 3 L of SC-His medium supplemented with 15% glucose after autoclaving. The reactor was set to 30°C, pH of 5.5, and a minimum dissolved oxygen of 40%. One blue LED panel (HQRP New Square 12” Grow Light Blue 517 LED 14W) was placed 20 cm from the cell culture. At this distance, the light panel output was 75 μmol/m^2^/s and covered ~7% of the available bulk surface area of the fermentation (Fig. 3a). The reactor was inoculated to an OD_600_ of 1 and the cells were grown in the dark (maintained by covering the reactor with black fabric) until the cultured reached an OD_600_ of 15, which took about 12 hours. At an OD_600_ of 15, the lights were turned on at 5s ON/ 95s OFF duty cycles and samples were taken 0 h, 3 h, and 24 h after light induction. At each time point, cell samples were diluted in ice-cold PBS to an OD_600_ of 0.5, kept on ice in the dark, and taken to flow cytometry.

### Screening of lactic acid producing strains

Starting from strain YEZ44, we integrated OptoAMP4 (EZ-L582) into the *HIS3* locus, creating YEZ336. Next, we transformed lactate dehydrogenase (LDH1) from *Lactobacillus casei* under P_GAL1-S_ (EZ-L605: 2μ plasmid with *URA3* selection) into YEZ366, creating YEZ423. Single colonies of YEZ423 were used to inoculate 1 mL of SC-Ura + 2% glucose media in 24-well plates and grown overnight at 30°C, 200 RPM, and under ambient light conditions. Each culture was then back-diluted into fresh media to OD_600_ = 0.1 and grown for 20 hours (until cultures reached an OD_600_ of 3) while grown in the dark (wrapped in aluminum foil). Cultures were then grown for 12 more hours under full light. Each culture was then centrifuged at 1,000 RPM for 5 minutes and cell pellets were resuspended in 1 mL of fresh SC-Ura + 2% glucose media. The plates were then sealed with Nunc Sealing Tape and incubated for 24 hours at 30°C under full blue light. Finally, cells were centrifuged, and supernatants were collected for HPLC analysis.

### Screening of isobutanol producing strains

We introduced enzymes of the mitochondrial isobutanol pathway – ketol-acid reducto-isomerase (*ILV5*), dihydroxyacid dehydratase (*ILV3*), CoxIV-tagged α-ketoacid decarboxylase (CoxIV-*ARO10*), and CoxIV-tagged alcohol dehydrogenase from *Lactococcus lactis* (CoxIV-*LladhA^RE1^*) – under constitutive promoters, along with acetolactate synthase (*ILV2*), the first enzyme in the mitochondrial isobutanol pathway, under P_GAL1-S_. These genes (EZ-L390: 2μ plasmid with *URA3* selection) were transformed into YEZ336, creating YEZ516.

Colonies of YEZ516 were used to inoculate 1 mL of SC-Ura + 2% glucose media in 24-well plates and grown overnight at 30°C, 200 RPM, and under ambient light conditions. Each culture was then back-diluted into new media to OD_600_ = 0.1 and grown for 20 hours (until cultures reached an OD_600_ of 3) and grown in the dark (wrapped in tinfoil). Cultures were then grown for 12 more hours under full light. Each culture was then centrifuged at 1,000 RPM for 5 minutes and cell pellets were resuspended in 1 mL of fresh SC-Ura + 2% glucose media. The plates were then sealed with Nunc Sealing Tape and incubated for 48 hours at 30°C and 200 RPM under 2s ON / 118s OFF pulsed light. Finally, cells were centrifuged, and supernatants were collected for HPLC analysis.

### Screening of naringenin producing strains

Starting from strain JCY125 (YEZ44 *ΔARO10)*, we integrated OptoAMP4 (EZ-L580) into the *HIS3* locus, creating YEZ480. We then integrated pMAL236, containing constitutively expressed enzymes to boost shikimate production^20^ – ribose-5-phosphate ketol-isomerase (*RKI1*), transaldolase (*TAL1*), transketolase (*TKL1*), pentafunctional aromatic enzyme (*ARO1*), feedback-insensitive 3-deoxy-D-arabino-heptulosonate-7-phosphate synthase (*ARO4^K229L^*), chorismate synthase (*ARO2*), and feedback-insensitive chorismite mutase (*ARO7^G141S^*) – into the *LEU2* locus of YEZ480 to create YEZ482. Next, we introduced constitutively expressed enzymes to overproduce tyrosine, phenylalanine, and malonyl-CoA^25,26^ – phenylalanine and tyrosine transaminase (*ARO8*), prephenate dehydrogenase (*TYR1*), prephenate dehydratase (*PHA2*), and phosphorylation inactivation-resistant acetyl-CoA carboxylase (*ACC1^S1157A^*) – through multi-copy integration of pMAL311 and pMAL399 into δ sites to make YEZ486. Finally, we introduced enzymes of the naringenin pathway – phenylalanine ammonium lyase from *Arabidopsis thaliana* (AtPAL2), a fusion of cinnamate 4-hydroxylase and NADPH-cytochrome P450 reductase from *A. thaliana* (AtC4H-AtATR2), 4-coumarate-CoA ligase from *A. thaliana* (At4CL2), naringenin-chalcone synthase *Hypericum androsaemum* (HaCHS), and chalcone isomerase from *Petunia hybrida* (PhCHI) – under constitutive promoters, along with tyrosine ammonia-lyase from *Flavobacterium johnsoniae* (FjTAL) and phenylalanine ammonia-lyase from *A. thaliana* (AtPAL2), the first enzymatic step towards naringenin biosynthesis from tyrosine and phenylalanine, under P_GAL1-S_. These genes were introduced using a 2μ plasmid (EZ-L645) with a *URA3* selection to transform YEZ486, creating YEZ488.

Colonies of YEZ488 were used to inoculate 1 mL of SC-Ura + 2% glucose media in 24-well plates and grown overnight at 30°C, 200 RPM, and under ambient light conditions. Each culture was then back-diluted into new media to OD_600_ = 0.1 and grown for 20 hours (until an OD_600_ of 3) in the dark (wrapped in aluminum foil). Cultures were then grown for 12 more hours under full light. Each culture was then centrifuged at 1,000 RPM for 5 minutes and cell pellets were resuspended in 1 mL of fresh SC-Ura + 2% glucose media. The plates were then sealed with Nunc Sealing Tape and incubated for 48 hours at 30°C and 200 RPM in full darkness. Finally, cells were centrifuged, and supernatants were collected for HPLC analysis.

### Three-phase fermentations

After screening 12 colonies for production of each chemical, we performed 3-phase fermentations (in quadruplicates) with the colony exhibiting the highest product yield, using different light schedules (full light, full darkness, or 1s ON / 79s OFF pulses) during the growth, and induction phases. For all fermentations, the growth phase was 20 hours following back-dilution of the overnight cultures; the induction phase was 12 hours immediately following the growth phase, and carried out in the same media; finally, the production phase initiated after centrifuging the cultures and resuspending the cells in fresh media. For lactic acid, the production phase was performed under full blue light for 24 hours. For isobutanol, the production phase was performed under 2s ON / 118s OFF light for 48 hours. For naringenin, the production phase was performed in full darkness for 48 hours. After the fermentation, the cultures were centrifuged and the supernatants collected for HPLC analysis.

### Quantification of Chemical Production

For lactic acid and isobutanol samples, plates were spun down at 1,000 RPM for 10 minutes to remove cells and other solid debris, and 300 μL of supernatant was taken for HPLC analysis. For naringenin samples, 0.6 mL of saturated cell cultures were mixed with 0.6 mL of methanol in 1.5 mL microcentrifuge tubes and incubated at 99°C for 5 minutes while vortexing every minute. Solutions were then spun down at 13,000 RPM for 30 minutes at 4°C in a benchtop micro-centrifuge (Eppendorf Centrifuge 5424) to remove cells and other solid debris, and 300 μL of supernatant was taken for HPLC analysis.

Concentrations of lactic acid, ethanol, isobutanol, and naringenin were quantified with high-performance liquid chromatography (HPLC), using an Agilent 1260 Infinity instrument (Agilent Technologies, Santa Clara, CA, USA). For lactic acid and isobutanol, samples were analyzed using an Aminex HPX-87H ion-exchange column (Bio-Rad, Richmond, CA, USA). The column was eluted with a mobile phase of 5 mM sulfuric acid at 55°C and a flow rate of 0.6 ml/min. Glucose, lactic acid, isobutanol, and ethanol were monitored with a refractive index detector (RID). For naringenin, samples were analyzed using Alltech Alltima C18 column (250 x 4.6 mm, 5 μm particle size) using 0.1% trifluoroacetic acid in acetonitrile (Solvent A), 0.1% trifluoroacetic acid in water (Solvent B) at 35°C, a flow rate of 0.9 ml/min, and the following gradient method: start at 10% Solvent A; from 0-4 min, linear increase of Solvent A from 10% to 70%; from 4-4.1 min, linear increase of Solvent A from 70% to 100%; hold at 100% Solvent A from 4.1-9 min; linear decrease of Solvent A from 100% to 10% A from 9-15 min; post-time run of 10 min. Naringenin was monitored with a diode array detector (DAD) using a detection wavelength of 290 nm. To determine analyte concentrations, peak areas were integrated and compared to those of standard solutions for quantification.

## Data Availability

The authors declare that all data supporting the findings of this study are available within the paper (and its supplementary information files), but original data that supports the findings are available from the corresponding authors upon reasonable request.

## Acknowledgements

We thank Dr. Christina DeCoste, Dr. Katherine Rittenbach, and the Princeton Molecular Biology Flow Cytometry Resource Center for assistance with flow cytometry experiments. We also thank the Avalos Lab members for useful discussions.

## Author contributions

E.M.Z. and J.L.A. conceived this project and designed the experiments. E.M.Z., M.A.L., J.-M. C., and P.O. constructed the strains and plasmids. E.M.Z. and M.A.L. executed the experiments. E.M.Z., M.A.L., J.E.T., and J.L.A. analyzed the data and wrote the paper.

