## Supplementary Information for "Optogenetic Amplification Circuits for Light-Induced Metabolic Control"

**Supplementary Sequences:**

**Sequence 1:** ***EL222 A79Q Mutant***

GGAGCAGATGATACTAGAGTTGAAGTCCAACCTCCAGCCCAATGGGTATTGGATTTG
ATCGAGGCCTCACCCATAGCAAGTGTTGTTAGTGATCCCAGATTGGCTGATAATCCT
TTGATTGCTATCAACCAGGCCTTTACCGATCTAACTGGGTATTCTGAGGAAGAATGT
GTTGGGCGTAATTGTAGGTTCTTACAGGGATCTGGTACTGAACCCTGGCTAACAGAT
AAGATCCGTCAAGGTGTTCGTGAGCATAAACCCGTGTTGGTTGAGATTTTGAATTAC
AAAAAGGATGGTACTCCTTTTCGTAATGCAGTTTTGGTGGCACCAATCTATGATGAC
GATGATGAATTACTATACTTCCTTGGTAGCCAAGTGGAAGTGGATGACGATCAACCG
AACATGGGCATGGCCCGTAGAGAACGTGCTGCGGAGATGTTAAAAACTCTTTCACC
ACGTCAATTAGAAGTCACAACTCTGGTTGCATCAGGCCTAAGAAATAAAGAAGTGG
CTGCCAGACTGGGTCTTTCAGAAAAAACTGTCAAAATGCACAGAGGTTTGGTAATG
GAAAAATTAAACTTAAAAACGAGTGCAGATTTAGTTAGAATTGCCGTAGAAGCAGG
TATTTAA

Highlighted is the A79Q mutation.

**Sequence 2: $P_{GAL10\text{-}M}$**

ATCGCTTCGCTGATTAATTAatcatcAAATAAGGCTAAAAACTAATCGCATTATCATCC
TATGGTTGTTAATTTGATTCGTTAATTTGAAGGTTTtaccaGCCAGGTTACTGCCAATTT
TTCCTCTTCATAACCATAAAAGCTAGTATTGTAGAATCTTTATTGTTCGGAGCAGTGC
GGCGCGAGGCACATCTGCGTTTCAGGAACGCGACCGGTGAAGACGAGGACGCACGG
AGGAGAGTCTTCCGTCGGAGGGCTGTCGCCCGCTCGGCGGCTTCTAATCCGTACTTC
AATATAGCAATGAGCAGTTAAGCGTATTACTGAAAGTTCCAAAGAGAAGGTTTTTTT
AGGCTAAGATAATGGGGCTCTTTACATTTCCACAACATATAAGTAAGATTAGATATG
GATATGTATATGGTGGTAATGCCATGTAATATGATTATTAAACTTCTTTGCGTCCATC
CAAAAAAAAAGTAAGAATTTTTGAAAATTCAATATAA

Highlighted are the removed Mig1 binding sites.

**Sequence 3: $P_{GAL1\text{-}M}$**

CGGATTAGAAGCCGCCGAGCGGGTGACAGCCCTCCGAAGGAAGACTCTCCTCCGTG
CGTCCTCGTCTTCACCGGTCGCGTTCCTGAAACGCAGATGTGCCTCGCGCCGCACTG
CTCCGAACAATAAAGATTCTACAATACTAGCTTTTATGGTTATGAAGAGGAAAAATT
GGCAGTAACCTGGTTGGTAAAACCTTCAAATGAACGAATCAAATTAACAACCATAG
GATGATAATGCGATTAGTTTTTAGCCTTATTTTAGTAGTAATTAATCAGCGAAGCG

ATGATTTTTGATCTATTAACAGATATATAAATGCAAAAACTGCATAACCACTTTAAC
TAATACTTTCAACATTTTCGGTTTGTATTACTTCTTATTCAAATGTAATAAAAGTATC
AACAAAAAATTGTTAATATACCTCTATACTTTAACGTCAAGGAGAAAAAACTATA

Highlighted are the removed Mig1 binding sites.

**Sequence 4: $P_{GAL1-S}$**

ccgggCGGATTAGAAGCCGCCGAGCGGGTGACAGCCCTCCGAAGGAAGACTCTCCTCC
GTGCGTCCTCGTCTTCACCGGTCGCGTTCCTGAAACGCAGATGTGCCTCGCGCCGCA
CTGCTCCGAACAATAAAGATTCTACAATACTAGCTTTTATGGTTATGAAGAGGAAAA
ATTGGATGATTTTTGATCTATTAACAGATATATAAATGCAAAAACGGATTAGAAGCC
GCCGAGCGGGTGACAGCCCTCCGAAGGAAGACTCTCCTCCGTGCGTCCTCGTCTTCA
CCGGTCGCGTTCCTGAAACGCAGATGTGCCTCGCGCCGCACTGCTCCGAACAATAAA
GATTCTACAATACTAGCTTTTATGGTTATGAAGAGGAAAAATTGGCAGTAACCTGGT
TGGTAAAACCTTCAAATGAACGAATCAAATTAACAACCATAGGATGATAATGCGAT
TAGTTTTTTAGCCTTATTTTAGTAGTAATTAATCAGCGAAGCGATGATTTTTGATCTA
TTAACAGATATATAAATGCAAAAACTGCATAACCACTTTAACTAATACTTTCAACAT
TTTCGGTTTGTATTACTTCTTATTCAAATGTAATAAAAGTATCAACAAAAAATTGTTA
ATATACCTCTATACTTTAACGTCAAGGAGAAAAAACTATAgcggccgcTAAAATC

Highlighted are extra binding sites from $P_{GAL1-M}$.

**Supplementary Tables:**

**Supplementary Table 1: Plasmids constructed for this study.**

| Plasmid | Position 1 | Position 2 | Position 3 | Position 4 | Position 5 | Position 6 | Position 7 | Marker | Vector type |
|---|---|---|---|---|---|---|---|---|---|
| EZ-L175 | $P_{PGK1}$_VP16-EL222_$T_{CYC1}$ | $P_{C120}$_*GAL4*_$T_{ACT1}$ | $P_{GAL1}$_*GFP*_$T_{ADH1}$ | EMPTY | EMPTY | EMPTY | EMPTY | *HIS*3 | Integration into *HIS*3 Locus |
| EZ-L316 | $P_{PGK1}$_VP16-EL222_$T_{CYC1}$ | $P_{C120}$_*GAL4*_$T_{ACT1}$ | $P_{GAL1\text{-}M}$_*GFP*_$T_{ADH1}$ | EMPTY | EMPTY | EMPTY | EMPTY | *HIS*3 | Integration into *HIS*3 Locus |
| EZ-L319 | $P_{PGK1}$_VP16-EL222_$T_{CYC1}$ | $P_{C120}$_*GAL4*_$T_{ACT1}$ | $P_{GAL10}$_*GFP*_$T_{ADH1}$ | EMPTY | EMPTY | EMPTY | EMPTY | *HIS*3 | Integration into *HIS*3 Locus |
| EZ-L320 | $P_{PGK1}$_VP16-EL222_$T_{CYC1}$ | $P_{C120}$_*GAL4*_$T_{ACT1}$ | $P_{GAL2}$_*GFP*_$T_{ADH1}$ | EMPTY | EMPTY | EMPTY | EMPTY | *HIS*3 | Integration into *HIS*3 Locus |
| EZ-L321 | $P_{PGK1}$_VP16-EL222_$T_{CYC1}$ | $P_{C120}$_*GAL4*_$T_{ACT1}$ | $P_{GAL7}$_*GFP*_$T_{ADH1}$ | EMPTY | EMPTY | EMPTY | EMPTY | *HIS*3 | Integration into *HIS*3 Locus |
| EZ-L339 | $P_{PGK1}$_VP16-EL222_$T_{CYC1}$ | $P_{C120}$_*GAL4*_$T_{ACT1}$ | $P_{GAL1\text{-}M}$_*GFP*_$T_{ADH1}$ | EMPTY | EMPTY | EMPTY | EMPTY | *HIS*3 | Integration into *HIS*3 Locus |
| EZ-L380 | $P_{PGK1}$_VP16-EL222_$T_{CYC1}$ | $P_{C120}$_*GAL4*_$T_{ACT1}$ | $P_{GAL10\text{-}M}$_*GFP*_$T_{ADH1}$ | EMPTY | EMPTY | EMPTY | EMPTY | *HIS*3 | Integration into *HIS*3 Locus |
| EZ-L390 | $P_{PGK1}$_*ILV3*_$T_{CYC1}$ | $P_{TEF1}$_CoxIV_ *adhA*$^{RE1}$_$T_{ACT1}$ | $P_{GAL1\text{-}M}$_*ILV2*_$T_{ADH1}$ | EMPTY | $P_{TEF1}$_*ILV5*_$T_{ACT1}$ | $P_{TDH3}$_CoxIV_*ARO10*_$T_{ADH1}$ | EMPTY | *URA3* | 2µ |
| EZ-L444 | $P_{PGK1}$_VP16-EL222_$T_{CYC1}$ | $P_{C120}$_*GAL4*_$T_{ACT1}$ | $P_{GAL1\text{-}S}$_*GFP*_$T_{ADH1}$ | EMPTY | EMPTY | EMPTY | EMPTY | *HIS*3 | Integration into *HIS*3 Locus |
| EZ-L545 | $P_{TEF1}$_VP16-EL222$^{A79Q}$_$T_{CYC1}$ | $P_{C120}$_*GFP*_$T_{ADH1}$ | EMPTY | EMPTY | EMPTY | EMPTY | EMPTY | *HIS*3 | Integration into *HIS*3 Locus |
| EZ-L560 | $P_{TEF1}$_VP16-EL222$^{A79Q}$_$T_{CYC1}$ | $P_{C120}$_*GAL4*_$T_{ACT1}$ | $P_{GAL1\text{-}S}$_*GFP*_$T_{ADH1}$ | EMPTY | EMPTY | EMPTY | EMPTY | *HIS*3 | Integration into *HIS*3 Locus |

| | | | | | | | | | |
|---|---|---|---|---|---|---|---|---|---|
| EZ-L580 | $P_{TEF1}$_VP16-EL222$^{A79Q}$_$T_{CYC1}$ | $P_{C120}$_*GAL4*_$T_{ACT1}$ | $P_{RNR2}$_*GAL80*_*PSD*_$T_{ADH1}$ | EMPTY | EMPTY | EMPTY | EMPTY | *HIS*3 | Integration into *HIS*3 Locus |
| EZ-L582 | $P_{TEF1}$_VP16-EL222$^{A79Q}$_$T_{CYC1}$ | $P_{RNR2}$_*GAL80*_*PSD*_$T_{ADH1}$ | EMPTY | EMPTY | $P_{C120}$_*GAL4*_$T_{ACT1}$ | $P_{GAL1-S}$_*GFP*_$T_{ADH1}$ | EMPTY | *HIS*3 | Integration into *HIS*3 Locus |
| EZ-L583 | $P_{TEF1}$_VP16-EL222$^{A79Q}$_$T_{CYC1}$ | $P_{ADH1}$_*GAL80*_*PSD*_$T_{ADH1}$ | EMPTY | EMPTY | $P_{C120}$_*GAL4*_$T_{ACT1}$ | $P_{GAL1-S}$_*GFP*_$T_{ADH1}$ | EMPTY | *HIS*3 | Integration into *HIS*3 Locus |
| EZ-L605 | $P_{GAL1-S}$_*LDH*_$T_{ADH1}$ | EMPTY | EMPTY | EMPTY | EMPTY | EMPTY | EMPTY | *URA3* | 2µ |
| EZ-L645 | $P_{TEF1}$_AtC4H-AtATR2_$T_{ACT1}$ | $P_{GAL1-S}$_AtPAL_$T_{ADH1}$ | $P_{PGK1}$_At4CL2_$T_{CYC1}$ | EMPTY | $P_{GAL1-S}$_FjTALL_$T_{TPS1}$ | $P_{TDH3}$_HaCHS_$T_{ADH1}$ | $P_{TEF1}$_PhCHI_$T_{ACT1}$ | *URA3* | 2µ |
| EZ-L891 | $P_{PGK1}$_VP16-EL222_$T_{CYC1}$ | $P_{C120}$_*GAL4*_$T_{ACT1}$ | $P_{GAL7-S}$_*GFP*_$T_{ADH1}$ | EMPTY | EMPTY | EMPTY | EMPTY | *HIS*3 | Integration into *HIS*3 Locus |
| pMAL236 | $P_{TDH3}$_RKI1_$T_{ADH1}$ | $P_{TEF1}$_TAL1_$T_{ACT1}$ | $P_{HHF2}$_TKL1_$T_{SA1}$ | $P_{CCW12}$_ARO4$^{K229L}$_$T_{ENO1}$ | $P_{PGK1}$_ARO7$^{G141S}$_$T_{CYC1}$ | $P_{TEF1}$_ARO2_$T_{ACT1}$ | $P_{TDH3}$_ARO1_$T_{ADH1}$ | *LEU2* | Integration into *LEU2* Locus |
| pMAL311 | $P_{PGK1}$_ARO8_$T_{CYC1}$ | $P_{TDH3}$_ACC1$^{S1157A}$_$T_{ADH1}$ | EMPTY | EMPTY | EMPTY | EMPTY | EMPTY | Zeocin | Integration into δ sites |
| pMAL399 | $P_{TEF1}$_TYR1_$T_{ACT1}$ | $P_{CCW12}$_PHA2_$T_{ENO1}$ | EMPTY | EMPTY | EMPTY | EMPTY | EMPTY | Zeocin | Integration into δ sites |

**Supplementary Table 2: Yeast Used for this Study**

| Strain Name | Genotype | Source |
|---|---|---|
| BY4741 | S288C *MAT***a** *his3Δ1 leu2Δ0 met15Δ0 ura3Δ0* | [1] |
| CEN.PK2-1C | *MAT***a** *his3Δ1 leu2*-3_112 *trp1*-289 *ura3*-53 | [2] |
| YEZ44 | CEN.PK2-1C *gal*80::KanMX *gal*4::HygB | [3] |
| YEZ72 | YEZ44 *HIS*$3_{cg}$::($P_{PGK1}$_VP16-EL222 _$T_{CYC1}$, $P_{C120}$_*GAL4*_$T_{ACT1,}$ $P_{GAL1}$_*GFP*_$T_{ADH1}$) | This Study |
| YEZ133 | YEZ44 *HIS*$3_{cg}$::($P_{PGK1}$_VP16-EL222 _$T_{CYC1}$, $P_{C120}$_*GAL4*_$T_{ACT1,}$ $P_{GAL1\text{-}M}$_*GFP*_$T_{ADH1}$) | This Study |
| YEZ139 | YEZ44 *HIS*$3_{cg}$::($P_{TEF1}$_VP16-EL222 _$T_{CYC1}$, $P_{C120}$_*GFP*_$T_{ADH1}$) | [3] |
| YEZ140 | CEN.PK2-1C *HIS3*$_{cg}$ | [3] |
| YEZ141 | YEZ44 *HIS*$3_{cg}$::($P_{PGK1}$_VP16-EL222 _$T_{CYC1}$, $P_{C120}$_*GAL4*_$T_{ACT1,}$ $P_{GAL10}$_*GFP*_$T_{ADH1}$) | This Study |
| YEZ142 | YEZ44 *HIS*$3_{cg}$::($P_{PGK1}$_VP16-EL222 _$T_{CYC1}$, $P_{C120}$_*GAL4*_$T_{ACT1,}$ $P_{GAL2}$_*GFP*_$T_{ADH1}$) | This Study |
| YEZ143 | YEZ44 *HIS*$3_{cg}$::($P_{PGK1}$_VP16-EL222 _$T_{CYC1}$, $P_{C120}$_*GAL4*_$T_{ACT1,}$ $P_{GAL7}$_*GFP*_$T_{ADH1}$) | This Study |
| YEZ163 | YEZ44 *HIS*$3_{cg}$::($P_{PGK1}$_VP16-EL222 _$T_{CYC1}$, $P_{C120}$_*GAL4*_$T_{ACT1,}$ $P_{GAL1\text{-}M}$_*GFP*_$T_{ADH1}$) | This Study |
| YEZ186 | CEN.PK2-1C *HIS3*::$P_{TEF1}$_GFP_$T_{CYC1}$ | [3] |
| YEZ189 | YEZ44 *HIS*$3_{cg}$::($P_{PGK1}$_VP16-EL222 _$T_{CYC1}$, $P_{C120}$_*GAL4*_$T_{ACT1,}$ $P_{GAL10\text{-}M}$_*GFP*_$T_{ADH1}$) | This Study |
| YEZ214 | YEZ44 *HIS*$3_{cg}$::($P_{PGK1}$_VP16-EL222 _$T_{CYC1}$, $P_{C120}$_*GAL4*_$T_{ACT1,}$ $P_{GAL1\text{-}S}$_*GFP*_$T_{ADH1}$) | This Study |

| | | |
|---|---|---|
| YEZ292 | YEZ44 *HIS*3$_{cg}$::(P$_{TEF1}$_VP16-EL222 $^{A79Q}$ _T$_{CYC1}$, P$_{C120}$_*GAL4*_T$_{ACT1,}$ P$_{GAL1-S}$_*GFP*_T$_{ADH1}$) | This Study |
| YEZ293 | YEZ44 *HIS*3$_{cg}$::(P$_{TEF1}$_VP16-EL222 $^{A79Q}$ _T$_{CYC1}$, P$_{C120}$_*GFP*_T$_{ADH1}$) | This Study |
| YEZ336 | YEZ44 *HIS*3$_{cg}$::(P$_{TEF1}$_VP16-EL222 $^{A79Q}$ _T$_{CYC1}$, P$_{RNR2}$_*GAL80_PSD*_T$_{ADH1}$, P$_{C120}$_*GAL4*_T$_{ACT1,}$ P$_{GAL1-S}$_*GFP*_T$_{ADH1}$) | This Study |
| YEZ337 | YEZ44 *HIS*3$_{cg}$::(P$_{TEF1}$_VP16-EL222 $^{A79Q}$ _T$_{CYC1}$, P$_{ADH1}$_*GAL80_PSD*_T$_{ADH1}$, P$_{C120}$_*GAL4*_T$_{ACT1,}$ P$_{GAL1-S}$_*GFP*_T$_{ADH1}$) | This Study |
| YEZ423 | YEZ336 + EZ-L605 | This Study |
| YEZ480 | JCy125 *HIS*3$_{cg}$::(P$_{TEF1}$_VP16-EL222 $^{A79Q}$ _T$_{CYC1}$, P$_{C120}$_*GAL4*_T$_{ACT1}$, P$_{RNR2}$_*GAL80_PSD*_T$_{ADH1}$) | |
| YEZ482 | YEZ480 *LEU2*::(P$_{TDH3}$_*RKI1*_T$_{ADH1}$, P$_{TEF1}$_*TAL1*_T$_{ACT1}$_P$_{PGK1}$_*PAD1*_T$_{CYC1}$, P$_{HHF2}$_*TKL1*_T$_{SSA1}$, P$_{CCW12}$_*ARO4*(K229L)_T$_{ENO1}$, P$_{TDH3}$_*ARO1*(D1407A)_T$_{ADH1}$_P$_{TEF1}$_*ARO2*_T$_{ACT1}$_P$_{PGK1}$_*ARO7*(G141S)_T$_{CYC1}$) | This Study |
| YEZ486 | YEZ482 YARCdelta5::( P$_{PGK1}$_*ARO8*_T$_{CYC1}$, P$_{TDH3}$_*ACC1*(S1157A)_T$_{ADH1}$, P$_{TEF1}$_*TYR1*_T$_{ACT1}$, P$_{CCW12}$_*PHA2*_T$_{ENO1}$) | This Study |
| YEZ488 | YEZ486 + EZ-L645 | This Study |
| YEZ516 | YEZ336 + EZ-L390 | This Study |
| JCY125 | CEN.PK2-1C *gal*80 *gal*4 aro10::KanMX | This Study |

**Supplementary Figures:**

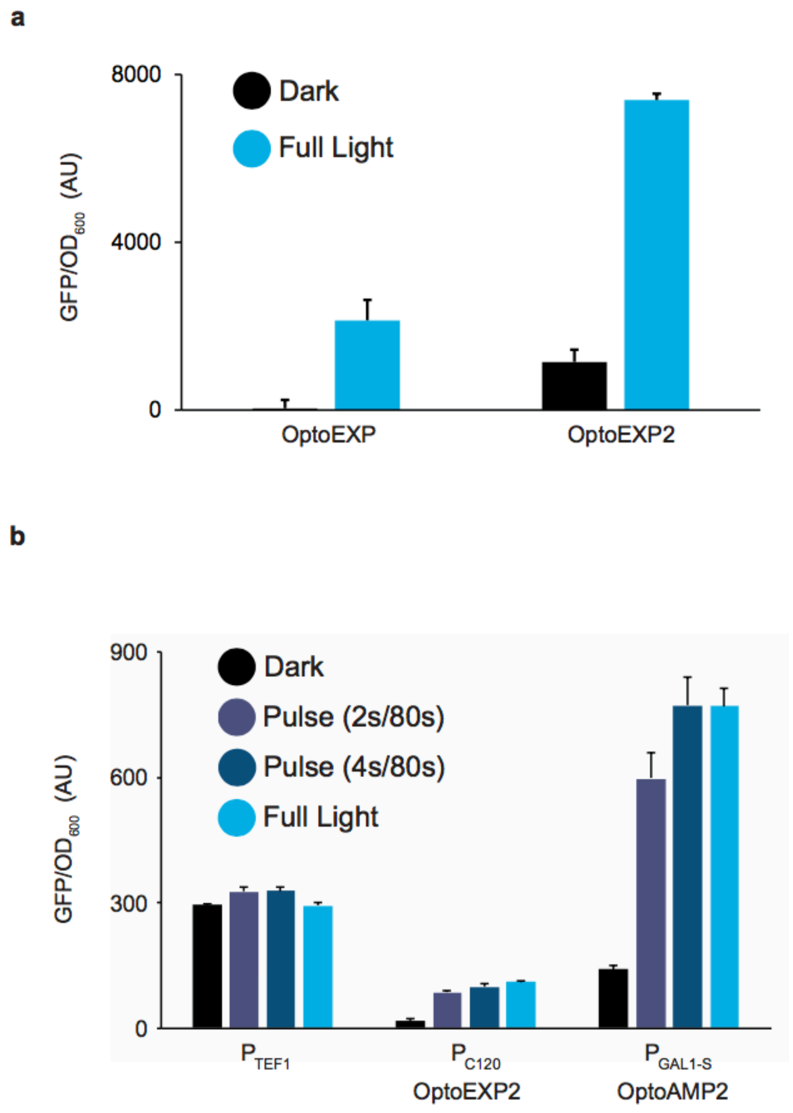


**Supplementary Figure 1. Characterization of OptoEXP2 and OptoAMP2**. (a) GFP expression using OptoEXP or OptoEXP2 in darkness or blue light. (b) GFP expression using OptoEXP2 (YEZ293) and OptoAMP2 with $P_{GAL1\text{-}S}$ (YEZ292) under different light conditions. All data are shown as mean values; error bars represent the s.d. of four biologically independent 1-ml sample replicates exposed to the same conditions.

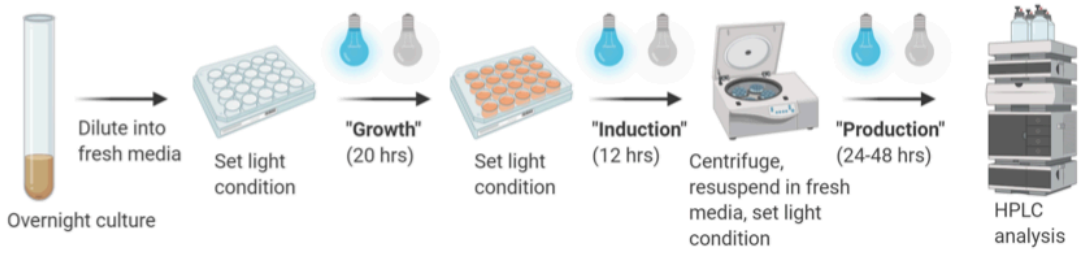


**Supplementary Figure 2. Three-phase fermentation protocol schematic.** Overview of fermentation setup. Cells are grown for 20 hours (Growth phase), induced for 12 hours (Induction phase), then centrifuged, resuspended in fresh media, and fermented for 24 hours (lactic acid) or 48 hours (isobutanol, naringenin) (Fermentation phase). Cultures are exposed to full light, intermediate light doses, or darkness during each phase. Created with Biorender.com.

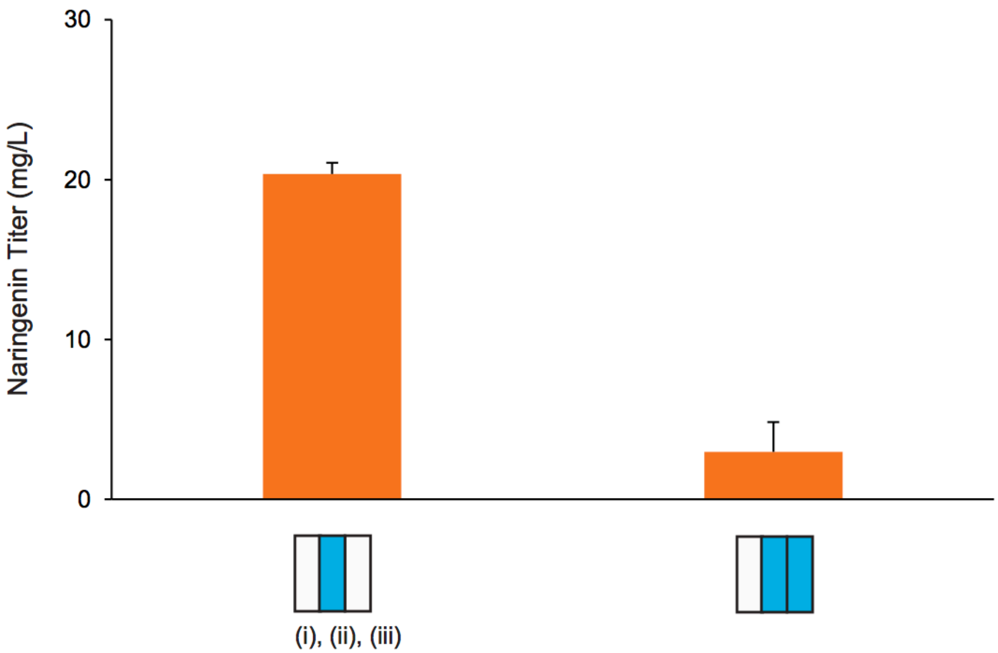


**Supplementary Figure 3. Naringenin titers under full light or darkness during the production phase.** Naringenin production using OptoAMP4 to control the expression of FjTAL and AtPAL2 from $P_{GAL1\text{-}S}$ (YEZ488) using a growth phase of full darkness, an induction phase of full light, and a production phase of darkness or full light. (i), (ii), and (iii) represent the light conditions used for the growth, induction, and production phases, respectively. All data are shown as mean values; error bars represent the s.d. of four biologically independent 1-ml sample replicates exposed to the same conditions.

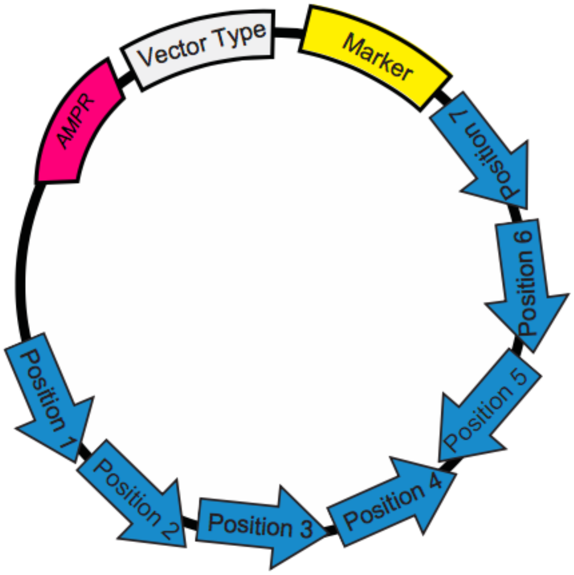


**Supplementary Figure 4. Scheme of vectors in this study.** General vector map that shows the relative orientation of the eight positions listed in Supplementary Table 1, in which different genes (including promoters and terminators) were assembled, using a previously described multiple gene insertion strategy[4]. All vectors have an ampicillin-resistance marker (*AmpR*) for cloning in *E. coli* and a selection marker for *S. cerevisiae* (Marker). Vector types include 2μ or integrative.

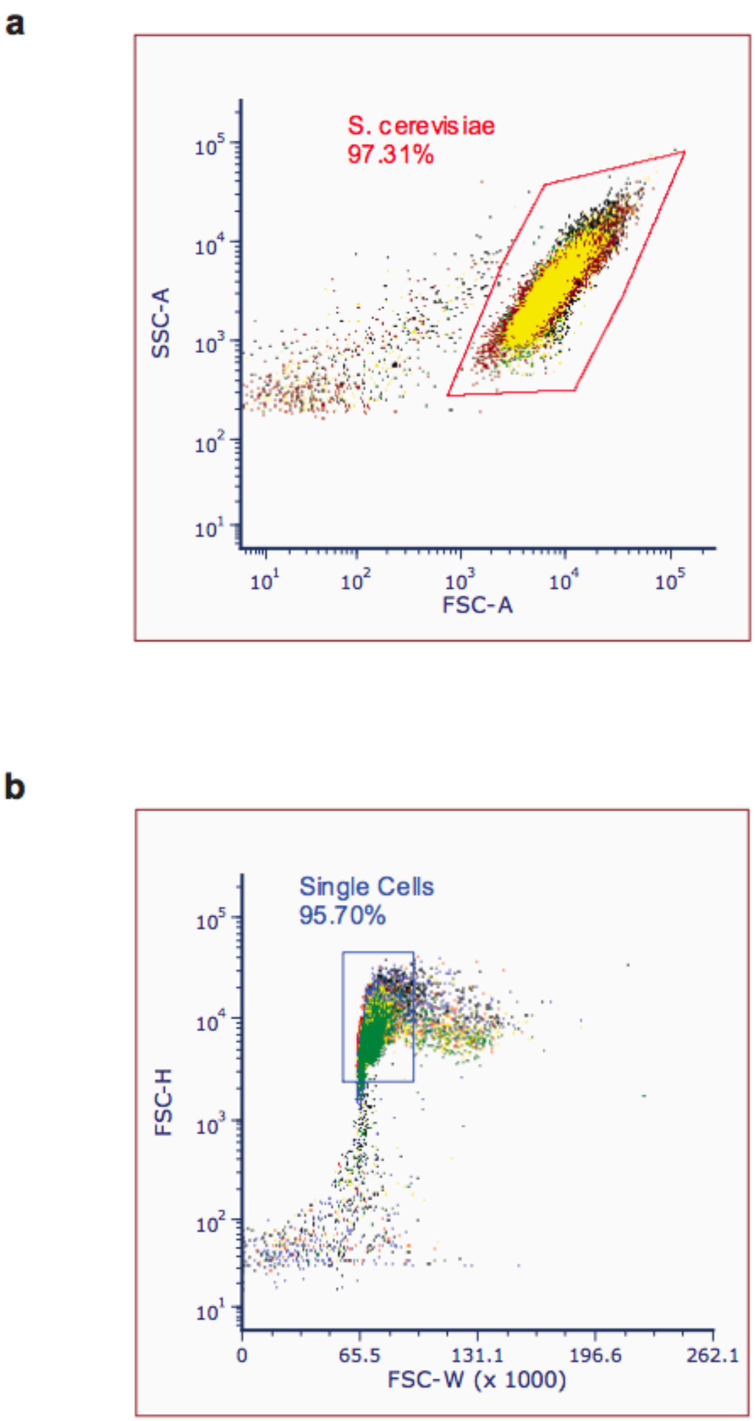


**Supplementary Figure 5. Flow cytometry gating strategy.** (a) Forward scatter area and side scatter area are used to distinguish yeast events from debris. (b) Forward scatter height and width are used to distinguish single-cell yeast events from doublets.
